# DONSON and FANCM associate with different replisomes distinguished by replication timing and chromatin domain

**DOI:** 10.1101/2020.03.19.999102

**Authors:** Jing Zhang, Marina A. Bellani, Ryan James, Durga Pokharel, Yongqing Zhang, John J. Reynolds, Gavin S. McNee, Andrew P. Jackson, Grant S. Stewart, Michael M. Seidman

## Abstract

Duplication of mammalian genomes requires replisomes to overcome numerous impediments during passage through open (eu) and condensed (hetero) chromatin. Typically, studies of replication stress characterize mixed populations of challenged and unchallenged replication forks, averaged across S phase, and model a single species of “stressed” replisome. However, in cells containing potent obstacles to replication, we find two different lesion proximal replisomes. One is bound by the DONSON protein and is more frequent in early S phase, in regions marked by euchromatin. The other interacts with the FANCM DNA translocase, is more prominent in late S phase, and favors heterochromatin. The two forms can also be detected in unstressed cells. CHIP-seq of DNA associated with DONSON or FANCM confirms the bias of the former towards regions that replicate early and the skew of the latter towards regions that replicate late.

## Introduction

Eukaryotic replisomes are multiprotein complexes consisting, minimally, of the CMG helicase [MCM2-7 (M), CDC45 (C), and GINS (go, ichi, ni, san) proteins (G)] which forms a ring around the leading strand template. Other components include the pol α, ε, and δ polymerases, MCM10, and a few accessory factors ^1–7^. The identification and characterization of the minimal components of biochemically active replisomes, the result of decades of extraordinary work from multiple laboratories, necessarily reflects studies with deproteinized model DNA substrates under carefully controlled conditions. However, *in vivo* there are hundreds of replisome associated proteins ^8–12^. Presumably this reflects the multiple layers of complexity that characterize replication of the genome in living cells. For example, three dimensional analyses of chromosome structure demonstrate two major domains. The A compartment contains euchromatin, which is accessible, transcriptionally active, and marked by specific histone modifications, such as H3K4me3. The B compartment, which is more condensed, contains inactive genes, many repeated elements, and is associated with different histone modifications, including H3K9me3 ^13^. In addition to the structural distinctions, regions of the genome are also subject to temporal control of replication during S phase. Sequences in Compartment A tend to replicate early in S phase, while those in B are duplicated in late S phase ^14,15^.

Other influential effectors of replisome composition are the frequent encounters with impediments, that stall or block either the progress of the CMG helicase or DNA synthesis. These include alternate DNA structures, protein: DNA adducts, DNA covalent modifications introduced by endogenous or endogenous reactants, depleted nucleotide precursor pools, etc. Replication stress activates the ATR (ATM- and Rad3-related) kinase, with hundreds of substrates, including MCM proteins ^16–18^, and stimulates the recruitment of numerous factors to stalled replication forks ^19–22^. These function in a variety of pathways to relieve obstacles, reconstruct broken forks, and restart replication.

We have developed an approach to studying replication stress imposed by an interstrand crosslink (ICL). While these have always been considered absolute blocks to any process requiring DNA unwinding ^23,24^, we found that replication could restart (traverse) past an intact ICL in the genome of living cells ^25^ (see also ^26^). Traverse of the ICL was dependent on ATR, and, in part, on the translocase activity of FANCM ^27,28^. FANCM was recruited to ICL proximal replisomes which were marked by phosphorylation of MCM2 by ATR. Furthermore, the association with FANCM was accompanied by remodeling of replisomes characterized by the loss of the GINS complex ^29^.

The partial dependence of ICL traverse on FANCM raised the question of what other factor(s) would support this activity. Recently, the DONSON protein was described as mutated in a microcephalic dwarfism syndrome ^30,31^. This essential protein, which has no recognizable structural features, associates with replisomes and contributes to the response to replication stress. In the work described here we find that, like FANCM, DONSON is complexed with ICL proximal replisomes also lacking the GINS proteins. The two “stressed” replisomes are distinguished by activity in different stages of S phase and different chromatin regions. In cells without ICLs DONSON and FANCM associate with sequences that show the same differential biases in replication timing and chromatin domain.

## Results

### DONSON contributes to ICL traverse

We have developed an approach to following replication in the vicinity of antigen tagged ICLs. Cells were treated with Digoxigenin-trimethylpsoralen and long wave ultraviolet light (Dig-TMP/UVA) and pulsed sequentially with CldU and IdU prior to spreading DNA fibers. Staining of the incorporated analogues and the Dig tag displays the outcomes of fork encounters with ICLs (**Fig. 1a**) (**Supplementary Fig. 1a, b**). Replication restart past ICLs (traverse) was reduced in cells deficient for FANCM, as shown previously ^25^. Recently, the DONSON protein was shown to contribute to the cellular response to replication stress ^30,31^. While DONSON does not appear to be a conventional DNA repair factor (it was not important for survival of cells exposed to cisplatin, **Supplementary Fig. 1c**), reduced expression of DONSON, either by siRNA knockdown (**Fig. 1b, Supplementary Fig. 1d**), or by mutation in patient derived cells (**Supplementary Fig. 1e**), did influence the results of the replication/fiber assay. Traverse frequency was reduced in these cells and declined further in doubly deficient cells, indicating that DONSON and FANCM were non epistatic for ICL traverse (**Fig. 1b**).

**1.**
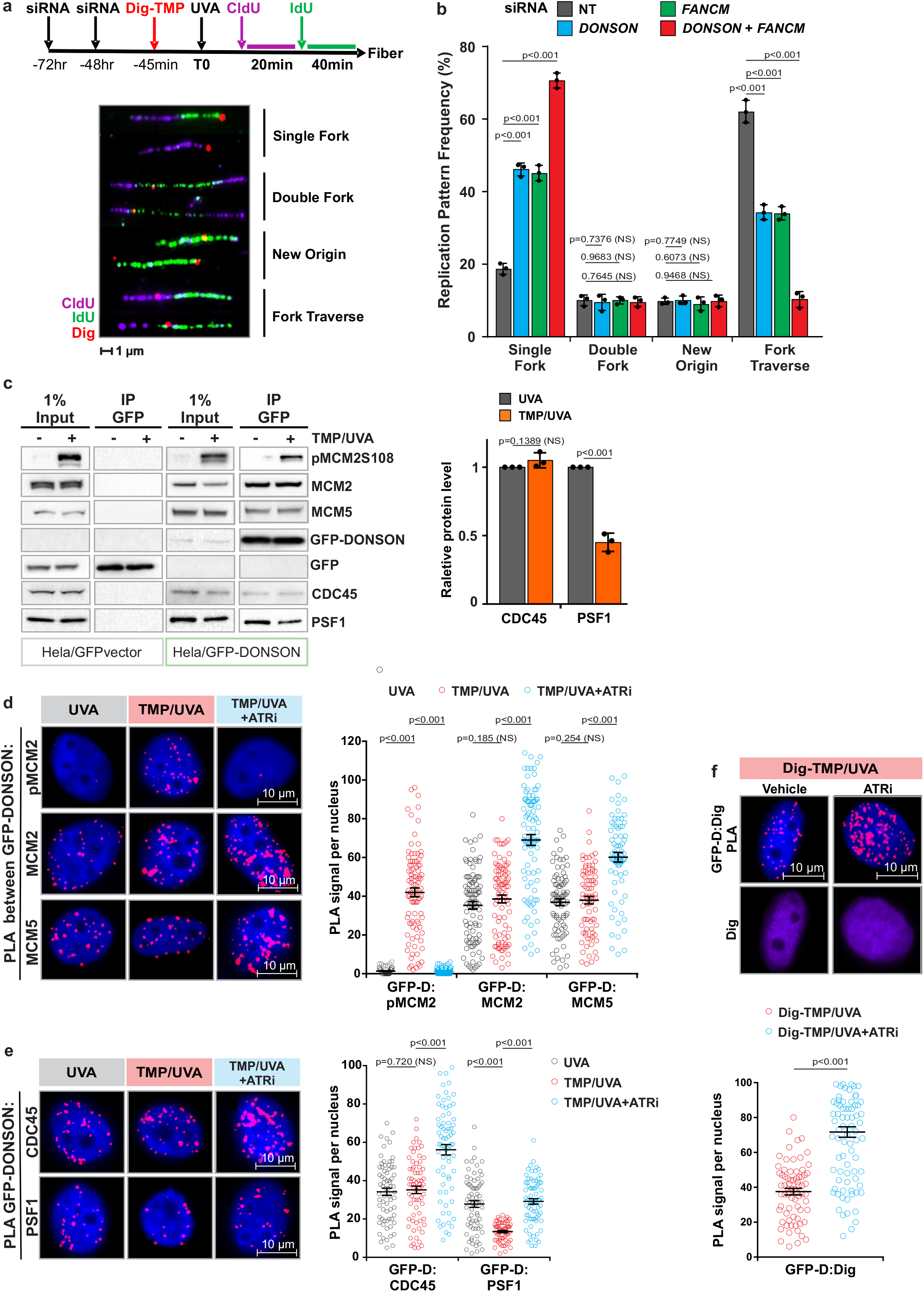
DONSON and FANCM operate in separate pathways to promote replication traverse. **a.** Schematic of the experimental procedure. HeLa cells were treated with siRNA against DONSON or FANCM or both. They were exposed to Dig-TMP/UVA and incubated with CldU and then IdU. Fibers were prepared and the patterns displayed by immunofluorescence against the analogues and immunoquantum detection (Q-dot 655, in red) for Dig tagged ICLs. Representative patterns are shown. **b**. Quantitation of pattern distribution from cells treated as indicated. Fibers with ICL encounters: NT= 417; siDONSON= 432; siFANCM= 417; siDONSON + siFANCM = 385, from 3 independent replicates. **c**. IP immunoblot of chromatin proteins from cells expressing GFP (panels 1, 2) or GFP-DONSON (panels 3, 4) exposed to UVA (−) or TMP/UVA (+). The identity of the proteins is indicated on the side. The amounts of PSF1 and CDC45 in the two samples were quantitated. Representative blot (n = 3) **d**. PLA test of the influence of ATR inhibition on GFP-DONSON interactions with pMCM2S108, MCM2, and MCM5. Number of nuclei: PLA between GFP-DONSON and pMCM2 in cells treated with UVA= 58, TMP/UVA= 94, TMP/UVA+ATRi= 55; PLA between GFP-DONSON and MCM2 in UVA= 95, TMP/UVA= 89, TMP/UVA+ATRi= 93; PLA between GFP-DONSON and MCM5 in UVA= 88, TMP/UVA= 79, TMP/UVA+ATRi= 73; from 3 biological replicates. **e**, PLA assessing the influence of ATR inhibition on GFP-DONSON interactions with CDC45 and PSF1. Scored nuclei of PLA between GFP-DONSON and CDC45 in UVA= 70, TMP/UVA= 71, TMP/UVA+ATRi= 73; Scored nuclei of PLA between GFP-DONSON and PSF1 in UVA= 71, TMP/UVA= 64, TMP/UVA+ATRi= 77; from 3 biological replicates. **f**. Influence of ATR inhibition on the PLA between GFP-DONSON and Dig tagged ICLs. Scored nuclei: Vehicle =72, ATRi = 87, 3 biological replicates. A two-sided unpaired t-test was used to calculate p-values for replication pattern frequency experiments and western blotting image analysis. Data are mean ± s.d. Mann-Whitney Rank sum test was used to calculate p-values for PLA experiments. Data are mean ± s.e.m. NS, not significant: p>0.05.

A relationship with replication and the replisome was indicated by co-immunoprecipitation of the endogenous DONSON protein or a GFP tagged DONSON with MCM proteins from untreated cells (no TMP/UVA) consistent with the prior report ^30^. DONSON was also complexed with CDC45 and the GINS proteins indicating association with the helicase functional form of the replisome (**Supplementary Fig. 1f**), in contrast to replisomes bound by FANCM ^29^. Proximity ligation assays (PLA) confirmed these interactions (**Supplementary Fig. 1g**). After TMP/UVA treatment, the association with MCM proteins and CDC45 was maintained while the interaction with PSF1, a GINS protein, was reduced (**Fig. 1c**). In the treated cells, PLA reported the proximity of DONSON and MCM proteins and also pMCM2S108, phosphorylated by ATR at S108 (**Fig. 1d**). The PLA between DONSON and PSF1 was positive in control cells and reduced in TMP/UVA cells (**Fig. 1e**) in agreement with the IP. These data demonstrated the association of DONSON with replisomes in cells with or without TMP/UVA treatment. Furthermore, they distinguished DONSON from FANCM, which, as shown previously, was not in complex with GINS proteins in either condition ^29^.

Inhibition of ATR blocks the association of FANCM with replisomes ^29^. In contrast, the PLA between DONSON and the ICLs was positive in control cells and increased after ATR inhibition (**Fig. 1f**). Thus, the response to ATR inhibition also differentiated the DONSON: replisome from the FANCM: replisome. These results are explained by a scenario in which encounters of DONSON: replisomes with ICLs are accompanied by the loss of GINS and traverse of the ICL. In the presence of the ATR inhibitor those replisomes accumulate at ICLs, traverse is blocked, and the GINS retained.

### DONSON and FANCM are on different replisomes

To determine if FANCM and DONSON were on the same replisomes we performed a sequential immunoprecipitation (IP) experiment (**Fig. 2a**). Chromatin was prepared from GFP-DONSON cells exposed to TMP/UVA, the DNA digested, and protein complexes incubated with antibody against PSF1, which served as a marker of a fully functional, “non stressed”, replisome. The precipitate contained the target PSF1, MCM2, DONSON, but neither FANCM nor pMCM2S108 (**Fig. 2b**). A second cycle of IP confirmed clearance of these replisomes **(Supplementary Fig. 2a)**. The supernatant was then incubated with antibody against GFP-DONSON. The IP contained GFP-DONSON and pMCM2S108, but no FANCM and, as expected, no PSF1. After another IP against GFP-DONSON, the remaining supernatant was incubated with antibody against FANCM. This IP contained FANCM, pMCM2S108, but no DONSON and no PSF1. Reversal of the order of the IP (FANCM before DONSON) did not change the results (**Supplementary Fig. 2b**). Thus, there were two DONSON associated replisomes: 1) replisome: CMG-D, independent of TMP/UVA, not marked by ATR phosphorylation, associated with the GINS; 2) replisome: CM-D, induced by TMP/UVA, with pMCM2S108 but not PSF1 or FANCM. The FANCM complex, replisome: CM-F, had pMCM2S108, but no GINS or DONSON. CDC45 and the auxiliary proteins MCM10, MCM8, and RAD51, were found in all samples (**Supplementary Fig. 2c**). These experiments were performed in HeLa cells expressing GFP-DONSON. In order to test the generality of these results we repeated the experiment in the hTERT immortalized diploid RPE1 cell line derived from retinal pigment epithelial cells and used in many studies of the cellular response to genotoxic stress ^32^. They displayed the same high frequency of ICL traverse as the HeLa and DONSON complemented patient derived cells (**Supplementary Fig. 2d**). The serial IP was performed except that antibody against the endogenous DONSON protein was employed. The results were identical to those with the GFP-DONSON HeLa cells **(Supplementary data Fig. 2e**).

**2.**
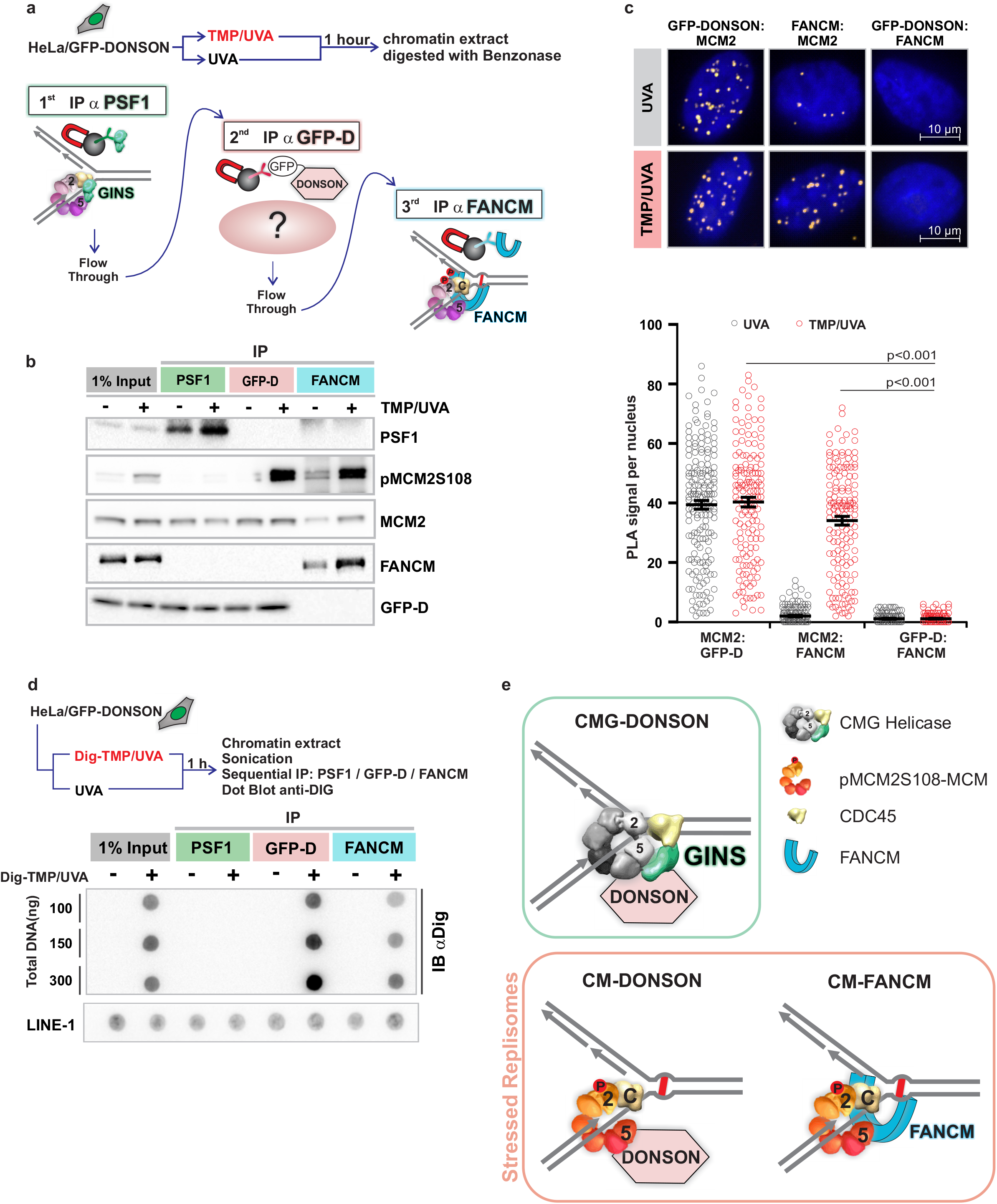
DONSON and FANCM are on different replisomes. **a.** Scheme of sequential IP against DONSON and FANCM associated replisomes. HeLa cells expressing GFP-DONSON were exposed to UVA only or TMP/UVA. Chromatin was prepared and digested with benzonase. This was followed by IP against PSF1 (to remove “unstressed” replisomes), then IP of the supernatant against GFP (to remove remaining DONSON associated proteins), and finally IP of the residual supernatant to capture FANCM bound proteins. **b**. Western blot analysis of sequential IP. **c**. PLA in cells exposed to UVA only or TMP/UVA shows interactions between GFP-DONSON and MCM2; and FANCM and MCM2; but not between GFP-DONSON and FANCM. Scored nuclei: PLA between GFP-D: MCM2, UVA treatment = 174; TMP/UVA =148; PLA between FANCM: MCM2, UVA = 142; TMP/UVA =145; PLA between GFP-D: FANCM, UVA =135; TMP/UVA =133. from 3 biological replicates. **d**. Association of replisomes with Dig-tagged ICLs. Chromatin was prepared from cells exposed to UVA or Dig-TMP/UVA and the DNA reduced to fragments of < 500 bp by sonication. Sequential IP was performed, and the DNA isolated from each fraction, dotted onto nitrocellulose and probed with an antibody to the Dig tag. LINE-1 repeat element served as a loading control. Representative blot (n = 2). **e**. Model summarizing the results of the sequential IP experiment. Mann-Whitney Rank sum test were used for analysis of PLA experiments. Data are mean ± s.e.m. NS, not significant: p>0.05.

PLA analyses with the GFP-DONSON HeLa cells agreed with the IP experiments. The interaction of DONSON with MCM2 in both UVA and TMP/UVA treated cells was positive (**Fig. 2c**). The PLA between FANCM and MCM2, which was detectable but low in cells without ICLs, was greatly increased in cells treated with TMP/UVA, while the PLA between DONSON and FANCM was negative in control and TMP/UVA treated cells.

We then tested the replisomes for association with ICLs. We treated cells with Dig-TMP/UVA and performed a sequential immunoprecipitation on chromatin sonicated to small DNA fragment size (sequential CHIP) in the order as in Fig. 2a. The DNA from each IP was recovered and examined for the presence of the Dig tag. There was no signal in the PSF1 sample, but both the subsequent precipitates were positive. Consequently, the ICLs were associated with replisomes containing CM-DONSON and CM-FANCM, but not CMG-DONSON (**Fig. 2d, e**, **Supplementary Fig. 2f)**.

### DONSON and FANCM replisomes at early and late S phase

To determine if CM-DONSON and CM-FANCM replisomes were in the same cell at the same time we performed sequential PLA on TMP/UVA treated cells grown on plates marked to facilitate re analysis of the same cells (**Fig. 3a, Supplementary Fig. 3a**). Images were taken of the DONSON: pMCM2S108 PLA, the plates were stripped of antibodies, followed by FANCM: pMCM2S108 PLA. The cells examined in the first analysis were re-imaged and the two images aligned in x, y, z (Methods). Some cells had more DONSON: pMCM2S108 signals than FANCM: pMCM2S108, while the opposite was true for others (**Fig. 3a**). Furthermore, although some cells had signals from both assays, they did not colocalize, indicating that these replisomes were in different genomic locations (**Supplementary Movie 1**).

**3.**
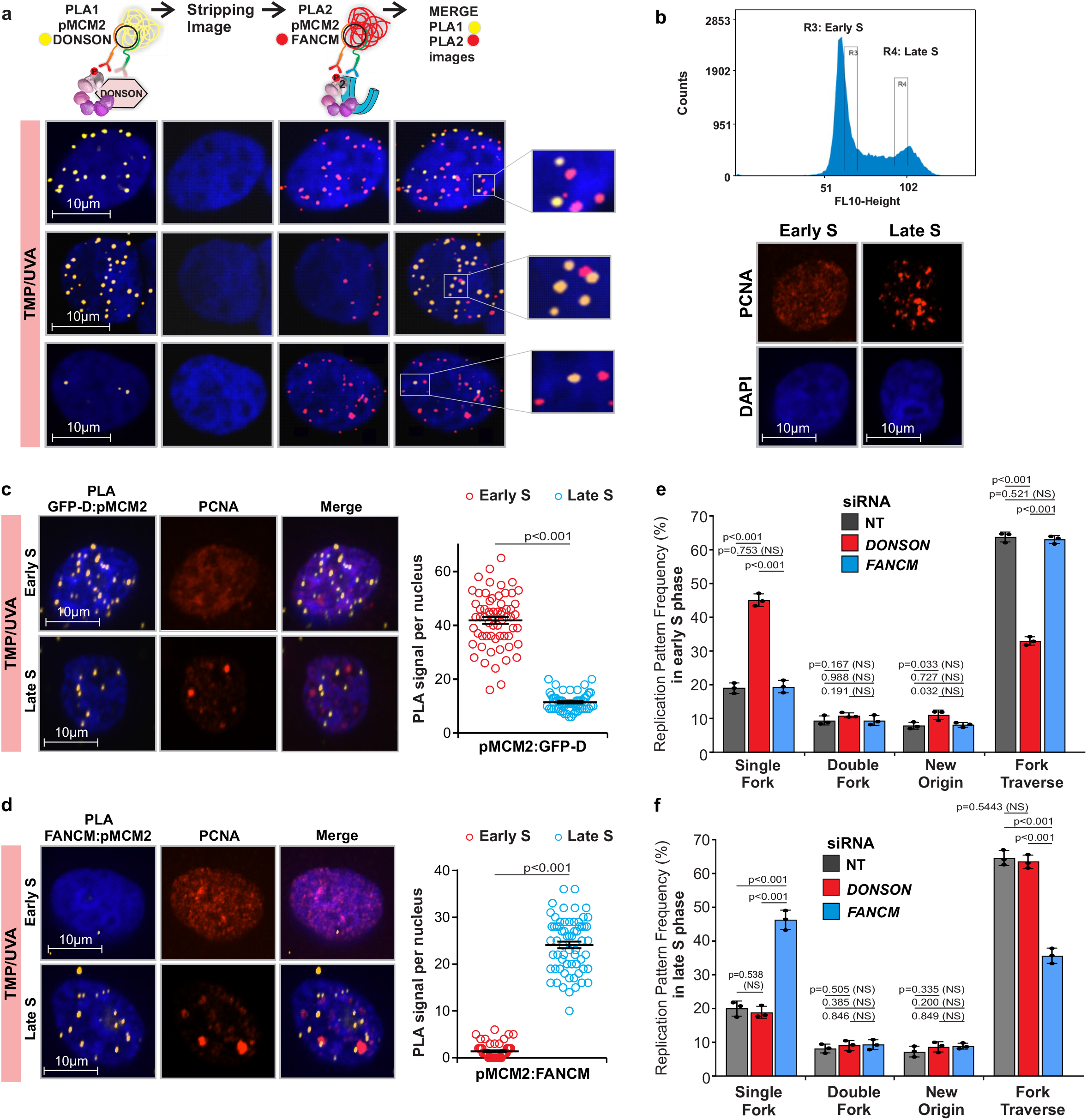
DONSON and FANCM replisomes are active in different stages of S phase. **a.** Analysis by sequential PLA of GFP-DONSON: pMCM2S108 complexes and then FANCM: pMCM2S108, in GFP-DONSON expressing cells exposed to TMP/UVA. After the first PLA the cells were photographed (first column of images) and the antibodies and PLA product stripped (second column). The second PLA was performed and the cells re-imaged (third column). The fourth column shows a merge of both images after image registration in the xyz planes using the DAPI signal. Shown are examples of cells with strong signals from both first and second PLA, or strong signals from the first but infrequent from the second, or weak from the first and strong from the second. The signals from the two PLA do not colocalize. **b**, Early and late S phase fractions were isolated from sorted cells. The PCNA staining pattern from each fraction. **c**. GFP-D: pMCM2S108 PLA in sorted early and late S phase cells. Scored nuclei: GFP-D: pMCM2S108 of early S phase= 62, late S phase= 60, from 3 biological replicates. **d**. FANCM: pMCM2S108 PLA in sorted early and late S phase cells. Scored nuclei: FANCM: pMCM2S108 of early S phase= 63, late S phase= 64, from 3 biological replicates. **e, f**. Influence of DONSON and FANCM on patterns of replication encounters with ICLs in early and late S phase cells. Cells were treated with siRNA against DONSON or FANCM, exposed to Dig-TMP/UVA and pulsed with nucleoside analogues as in Fig 1a. Cells were sorted, and fiber patterns from early and late S phase analyzed. Mann-Whitney Rank sum test were used for analysis of PLA experiments. Data are mean ± s.e.m. NS, not significant: p>0.05.

In an effort to understand the basis of these results, we treated cells with TMP/UVA and then recovered early and late S phase cells by flow cytometry (**Fig. 3b, Supplementary Fig. 3b**). The PLA between the Dig-tagged ICLs and pS108MCM2 showed equal frequencies of ICL proximal stressed replisomes in the two cell fractions (**Supplementary Fig. 3c**). DONSON: pMCM2S108 and FANCM: pMCM2S108 PLAs were performed on each group. The DONSON complex was about 4-fold more frequent in early S phase than in late, while the FANCM complex was about 10-fold more frequent in late S phase than in early (**Fig. 3c, d**). The negative PLA for both partner sets in G_1_ phase cells provided an important internal control for the specificity of the reagents and assay (**Supplementary Fig. 3d**).

The clear distinction between the early and late S phase fractions reflected the separation of the early S phase cells from those in late S phase. On the other hand, when we examined mid S phase cells the pronounced difference between the PLA frequencies of the two stressed replisomes was lost, indicating that both stressed replisomes were present in mid S phase cells (**Supplementary Fig. 3e**).

The influence of DONSON on replication fork encounters with ICLs in the early and late stages of S phase was tested in cells treated with siRNA/DONSON. There was an increase in single fork stalling events and a decline in traverse frequency in the early S phase cells, while there was little change in late S phase cells (**Fig. 3e, f**). Conversely, in cells treated with siRNA/FANCM there was an increase in single fork stalling and a decline in traverse frequency in late S phase cells, with relatively little effect on early S phase patterns (**Fig. 3e, f**). Thus, the DONSON: stressed replisome made a greater contribution to the traverse patterns in early S phase than in late, while the FANCM: replisome was more important in late S phase. Consequently, deficiencies in one or the other would differentially influence the outcome of replisome encounters with ICLs depending on the stage of S phase.

Alu sequences are replicated in early S phase, Satellite 3 sequences are replicated in late S phase, while LINE-1 elements are replicated throughout ^33^. Cells were treated with TMP/UVA and DNA isolated from each fraction from the sequential CHIP (as in Fig. 2f) and examined for the presence of these repeats. As expected, the replisome marked by PSF1 was associated with all the sequences. However, the recovery of Alu sequences was biased towards the replisome: CM-DONSON fraction, while the recovery of Satellite 3 was greater with the replisome: CM-FANCM (**Fig. 4a**). LINE-1, which replicates throughout S phase, was found in all fractions.

**4.**
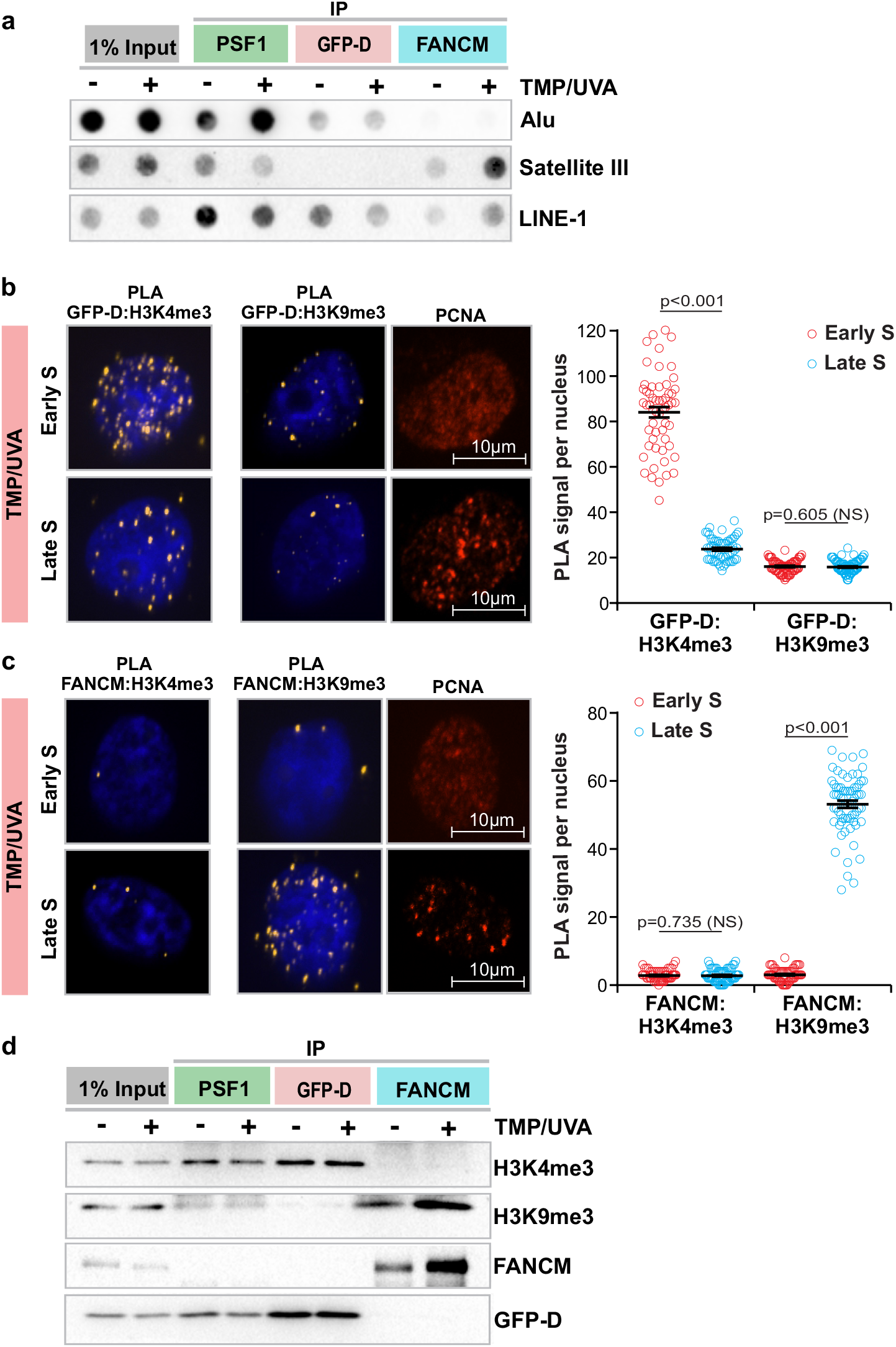
Relationship of DONSON and FANCM to replication timing and chromatin domain. Cells were treated with either UVA or TMP/UVA. **a**. Sequential IP demonstrates association of early replicating Alu sequences with DONSON and late replicating Satellite 3 sequences with FANCM. LINE-1 elements replicate throughout S phase and are found in all fractions. **b**. DONSON interaction with the H3K4me3 euchromatin mark is more frequent in early S phase cells than in late S phase, while there is little interaction with the H3K9me3 heterochromatin mark in either stage. Sorted early and late S phase cells were examined by PLA. Scored nuclei: PLA between GFP-D: H3K4me3 of early S phase= 67, late S phase= 64, PLA between GFP-D: H3K9me3 of early S phase= 70, late S phase= 85, from 3 biological replicates. **c**. FANCM interaction with H3K9 me3 heterochromatin mark is biased towards late S phase, while there is low interaction frequency with H3K4me3 in either stage. Scored nuclei: PLA between FANCM: H3K4me3 of early S phase= 64, late S phase= 66, PLA between FANCM: H3K9me3 of early S phase= 67, late S phase= 77, from 3 biological replicates. **d**. Sequential IP demonstrates greater association of DONSON with H3K4me3 than H3K9me3 and greater association of FANCM with H3K9me3 than H3K4me3. Mann-Whitney Rank sum test were used for analysis of PLA experiments. Data are mean ± s.e.m. NS, not significant: p>0.05.

Active genes replicate in early S phase, and are found in euchromatin, marked by histone H3K4 trimethylation ^34^, while inactive genes replicate late, and are in heterochromatin, characterized by H3K9 trimethylation ^35^. We treated cells with TMP/UVA and examined the proximity of GFP-DONSON to the two chromatin marks in early and late S phase cells. The PLA with H3K4me3 showed a 4-fold higher signal frequency in early S phase than in late, while the PLA with H3K9me3 was much weaker in both stages (**Fig. 4b**). The PLA between FANCM and H3K4me3 was quite low in both early and late S phase cells, while the signal with H3K9me3 was about 10-fold stronger in late S phase than in early S phase cells (**Fig. 4c**). These experiments were repeated in RPE1 cells with identical results (**Supplementary Fig. 4a, b).** They were also confirmed by sequential chromatin IP in both cell lines which showed that H3K4me3 was associated with the replisome: CM-DONSON complex, while the replisome: CM-FANCM was associated with H3K9me3 (**Fig. 4d, Supplementary Fig. 4c**). The results of these experiments confirmed the appearance of replisomes differing by association with either FANCM or DONSON in cells exposed to replication stress imposed by the ICLs.

### DONSON and FANCM replisomes in untreated cells

The preceding experiments characterized replisomes in cells containing ICLs and demonstrated the bias of DONSON replisomes towards early S phase. Previously, DONSON was shown to be bound to replisomes in cells without exposure to a DNA reactive compound ^30^, leaving open the question of whether it was complexed with all replisomes, or only a subset. To address this chromatin proteins from untreated cells were subjected to sequential IP, first with DONSON as the target, after which the supernatant was incubated with antibody against PSF1 to recover remaining functional replisomes. Two complexes were recovered: replisome: CMG-DONSON and, subsequently, replisome: CMG (**Fig. 5a**). These results identified two forms of the replisome in unstressed cells: one with DONSON and one without. We then asked if DONSON replisomes in untreated cells were more or less abundant in different stages of S phase. The PLA between GFP-DONSON and PSF1 showed about a 3.5 fold bias towards early S phase (**Fig. 5b**). The proximity of FANCM to MCM2, albeit at quite low frequency (Fig 2c, **Fig. 5c**), was biased to late S phase in non-treated cells (**Fig. 5c**). The low frequency interaction of FANCM with replisome proteins was also observed by immunoprecipitation (**Fig. 5d**).

**5.**
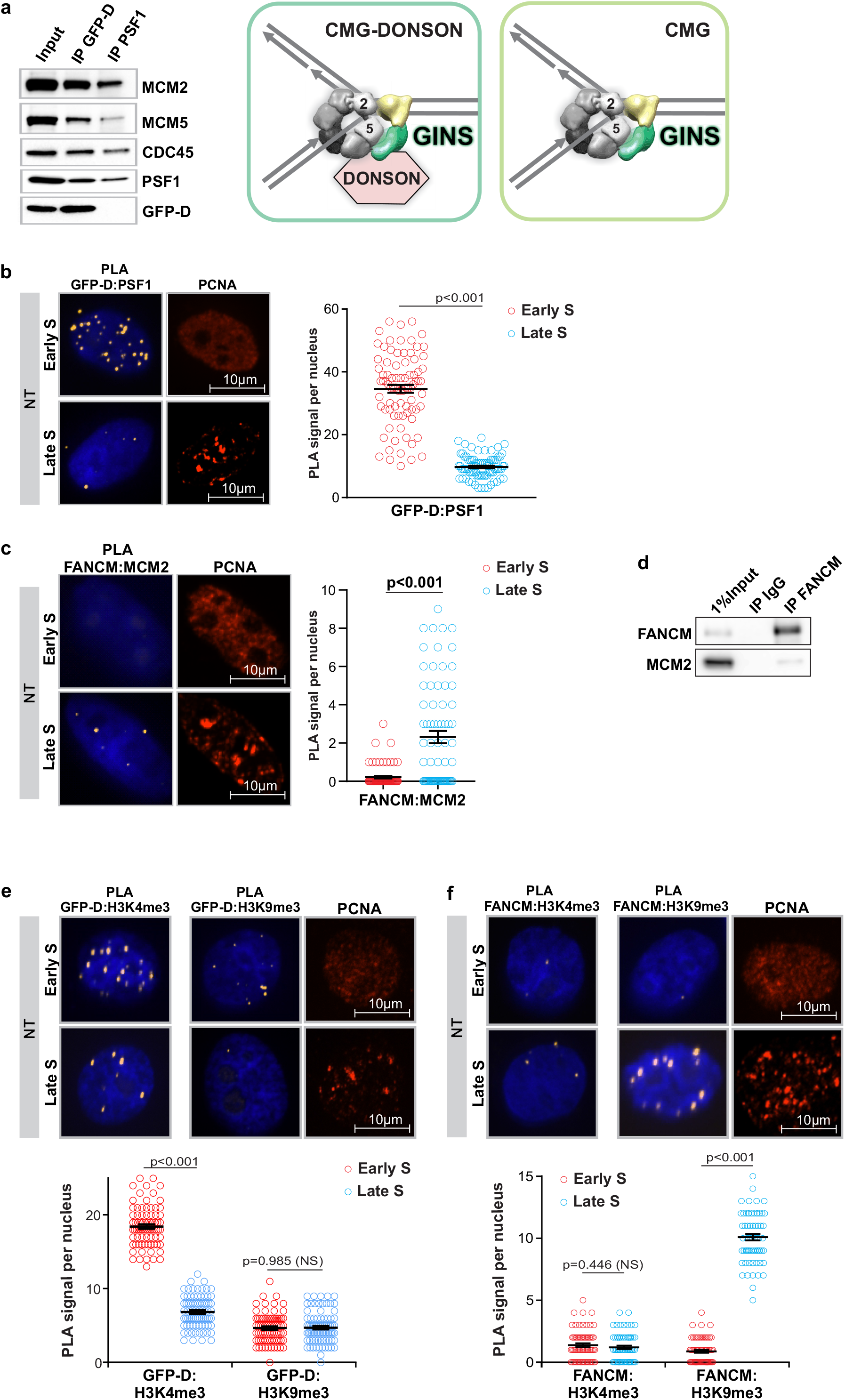
Interactions of DONSON and FANCM with replisomes and chromatin in non-treated cells. **a**. DONSON associates with some, but not all, replisomes in untreated cells. Chromatin was prepared from untreated GFP-DONSON-HeLa cells and sequential IP performed, first against GFP-DONSON, and then against the GINS protein PSF1 from the residual supernatant. **b**. PLA of GFP-DONSON and PSF1 demonstrates DONSON associated replisomes are more frequent in early S phase than in late S phase in NT cells. Scored nuclei: PLA between GFP-D: PSF1 early S phase= 82, late S phase= 81, from 3 biological replicates. **c**. PLA between FANCM and MCM2 demonstrates low level of FANCM associated replisomes in late S phase in non-treated cells. Scored nuclei: PLA between FANCM: MCM2 of early S phase= 73, late S phase= 75, from 3 biological replicates. **d.** IP of FANCM demonstrates low level interaction with replisome protein MCM2. **e**. PLA between GFP-DONSON and H3K4me3 or H3K9me3. Scored nuclei of GFP-DONSON and H3K4me3 in early S phase= 79, late S phase= 78; Scored nuclei of GFP-DONSON and H3K9me3 in early S phase= 77, late S phase= 82, from 3 biological replicates. **f**. PLA between FANCM and H3K4me3 or H3K9me3. Scored nuclei of FANCM and H3K4me3 in early S phase= 64, late S phase= 65; Scored nuclei of FANCM and H3K9me3 in early S phase= 68, late S phase= 65, from 3 biological replicates. Mann-Whitney Rank sum test was used to calculate p-values for PLA experiments. Data are mean ± s.e.m. NS: not significant (p>0.05).

We also asked about the proximity of DONSON and FANCM to modified histones in untreated cells. Cells were sorted and examined by PLA between GFP-DONSON or FANCM and H3K4me3 or H3K9me3. The DONSON: H3K4me3 signals were distributed throughout the nuclei and were about 3-fold more frequent in early S phase than in late, while there was little signal with H3K9me3 in either stage (**Fig. 5e**). There was minimal association between FANCM and H3K4me3 in either stage, while the interaction with H3K9me3 was weak in early S phase but about 10-fold stronger in late S phase (**Fig. 5f**). The FANCM: H3K9me3 PLA signals were largely localized on the nuclear periphery, reflecting the association of H3K9me3 chromatin with nuclear lamina ^36^. Thus, DONSON and FANCM were largely resident in different chromatin domains without requirement for ICL induced replication stress. Similar results were acquired with RPE1 cells (Supplementary **Fig. 5a, b**).

### Association of DONSON and FANCM with genomic sequences

The bias in replication timing and genome location indicated by the preceding experiments with untreated cells prompted a CHIP-seq analysis of DONSON and FANCM associated DNA in cells without ICLs (see Discussion). Chromatin was prepared, sonicated, and immunoprecipitated against GFP-DONSON or FANCM. DNA was isolated and subjected to Next Gen sequence analysis. The enrichment of FANCM and GFP-DONSON [log2(ChIP/input)] across individual chromosomes was compared to data on replication timing, and the Hi-C compartments A and B. The distribution of DONSON and FANCM associated sequences in most regions in chromosomes such as 1, 5, 9 matched well with the replication timing and Hi-C A and B compartments, respectively (**Fig. 6a, Supplementary Fig. 6a, c**). Chromosomes such as 6, 10, 12, showed little overlap between DONSON and FANCM, but the correlations with early and late replicating DNA and the A and B compartments were not as strong (**Supplementary Fig. 6b, d, e**). Additionally, there were chromosomes (14, 15) in which the DONSON and FANCM signals were intermingled (**Supplementary Fig. 6f, g**). There was no correspondence between the regions associated with DONSON or FANCM and fragile sites in any chromosome. Violin plots of DONSON and FANCM associated DNA sequences (across the entire genome) that were enriched relative to the input were skewed towards sequences that were early replicating and in compartment A or late replicating and in compartment B, respectively (**Fig. 6b**).

**6.**
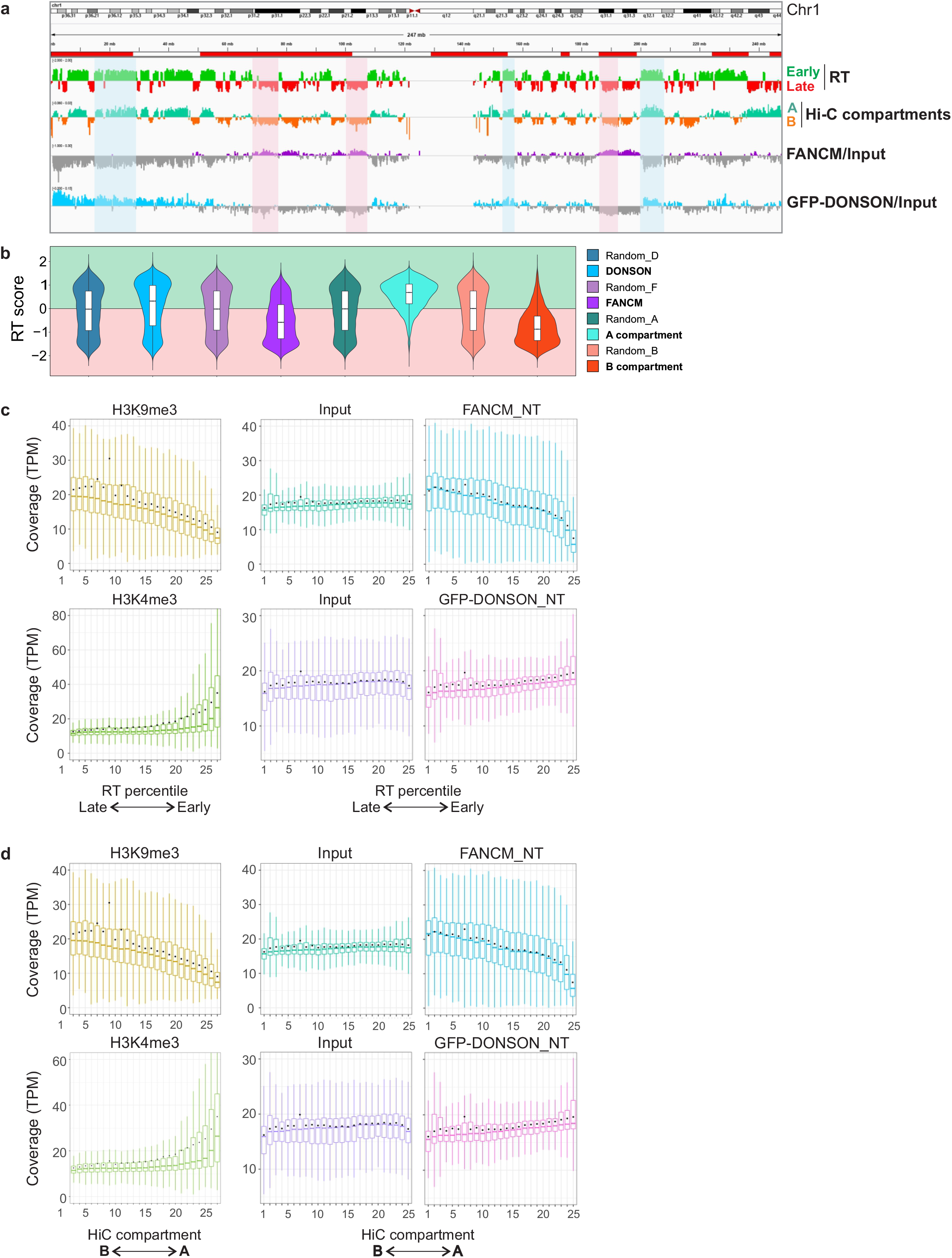
ChIP-Seq Analysis of genome-wide distribution of FANCM and GFP-DONSON. **a.** Representative profile from chromosome 1, comparing RT (RT = log2(Early/Late)) in Hela cells, A/B compartments as defined by the eigenvector calculated from Hi-C data from Hela cells, and FANCM and GFP-DONSON distribution (enrichment = log2(ChIP/input)) in Hela cells stably expressing GFP-DONSON. Shadowed in red are some examples of late replicating regions aligning with FANCM enriched genomic regions. In blue are highlighted some early replicating regions showing correspondence with GFP-DONSON enriched regions. In the RT profile, positive and negative values correspond to early and late replication respectively. In the eigenvector profile, they correspond to the A and B compartments. Regions containing fragile sites are marked by red bars above the profiles. **b.** Violin plot displaying the distribution of replication timing of 50 Kb genomic windows enriched in FANCM or GFP-DONSON ChIP (log2[ChIP/Input] > 0) and the A and B Hi-C compartments (eigenvector >0, <0, respectively), each compared to a matching number of randomly selected genomic windows of the same size. **c.** Coverage of H3K9me3, H3K4me3, FANCM and GFP-DONSON of 50 Kb genomic windows within 25 replication timing quantiles, going from late to early replicating regions, in Hela cells expressing GFP-DONSON. **d.** Coverage of H3K9me3, H3K4me3, FANCM and GFP-DONSON of 50 kb genomic windows within 25 eigenvector quantiles, going from B to A Hi-C compartments in Hela cells expressing GFP-DONSON.

In order to evaluate the relationship between DONSON and FANCM across the entire genome and early or late replicating loci, we calculated the coverage of the respective ChIP-seq results in replication timing quantiles in the cells. As proof of principle, we also calculated the coverage of H3K9me3 and H3K4me3 histone marks, using published data (Methods). As expected, the permissive chromatin mark H3K4me3 was progressively enriched towards early replicating regions of the genome, while the repressive histone mark H3K9me3 was progressively enriched towards late replicating regions. FANCM Chip-seq data were increasingly enriched towards late replicating regions, similar to H3K9me3 (**Fig. 6c**). The DONSON Chip-seq showed a bias towards the quantiles that associated with early replication, although it was not as pronounced as the FANCM linkage to late replication. Another comparison was to the continuum of A – B chromatin compartments defined by Hi-C. Again, there was a clear bias in the sequences captured by FANCM towards the B compartment associated with silent chromatin and late replicating sequences. DONSON bound sequences were weighted towards the A compartment, but not as strongly as H3K4me3 (**Fig. 6d**).

## Discussion

In living cells many more proteins associate with replisomes than are required for “minimal” biochemical reconstructions ^37,12^. These interactions may be constitutive or induced by replication stress, but are typically interpreted as representing a single complex (see Introduction) ^15,22,38–42^. An alternative view, that there are multiple, distinguishable, replisome variants, either constitutive or in response to stress, has received less attention. Our results demonstrate two compositionally different replisomes in “unstressed” cells, and an additional two in cells containing potent blocks to replication. Furthermore, we find that the different replisomes are also distinguished by replication timing and chromatin location.

DONSON bound replisomes were constitutively more prevalent in genomic regions with euchromatin histone marks, in chromatin compartment A, and were associated with early replicating elements. FANCM was more frequent in heterochromatin, in compartment B, and biased towards late replicating regions. The clarity of the data supporting these conclusions was dependent on experiments in which cells from well separated stages of S phase were analyzed (the PLA experiments), or chromatin complexes containing DONSON were separated from those bound by FANCM (the sequential IP). However, for practical reasons the CHIP-seq analyses were with unsorted cells which necessarily included cells from all stages of S phase, blurring the distinction between early and late stages (see Fig. S3e). Nonetheless, the CHIP-seq data, summed over the entire genome, were in accord with the conclusions of the experiments with early and late replicating cells. The examination of the patterns from individual chromosomes revealed some in excellent agreement with the early/late bias of DONSON/FANCM, while the results with others were not as clear. Generally, the distinctions were stronger for the FANCM bound sequences than for those of DONSON, in agreement with the results from the PLA experiments. It should be noted that at best these measurements will reflect the location of DONSON and FANCM in regions of the genome rather than at specific sites defined by several nucleotides, as would be the case with transcription factors. Factors that are involved in DNA transactions that function throughout the entire genome, or in enormous domains such as those assigned to the A and B compartments, are unlikely to be present at the same location at the same time across a cell population. These considerations have also been noted in analyses of the relationship between chromatin folding compartments and replication timing ^15,43^.

The presence, in active chromatin, of constitutive DONSON replisomes (replisome: CMG-D) suggests a cellular anticipation of replication stress in regions that are more susceptible to DNA damage and collisions with transcriptional R loops ^44–46^. Thus defects in DONSON ^30,31^ would preferentially influence the response to replication stress in active gene regions of the genome. DONSON is mutant in a microcephalic dwarfism syndrome. Inefficiencies in transit through transcriptionally active gene regions ^47^ could have adverse effects on completion of S phase and consequently, cell number, resulting in the smaller brain and body size that are features of individuals with DONSON mutations.

On the other hand, our results indicate that in unstressed cells the association of FANCM with replisomes is infrequent. It is possible that the FANCM: replisomes in untreated cells result from encounters of replisomes with endogenous blocks. Our data demonstrate the bias of FANCM to regions that replicate late and are marked by histone modifications consistent with heterochromatin. Consequently, we suggest that replisomes that encounter blocks in these domains are in environments with associated FANCM, and ready targets for FANCM recruitment. These regions contain “difficult to replicate” sequences ^35^. FANCM, which is an ancient protein with equivalents in archaea ^27^, may have evolved, in part, to respond to replication blocks in sequences with a propensity to stall replication. In FANCM deficiency disorders ^48,49^ we would anticipate that the fault in at least a component of the response to replication stress would be in heterochromatin ^50^.

In previous work we showed that the FANCM increment of ICL traverse was dependent on the translocase activity, while the loss of GINS required only the association of the protein, including a translocase inactive version, with the replisome complex ^29^. Thus, the functions of the protein in the traverse assay could be separated into at least two steps. We suggest that the replication restart pathway is multi step, requiring an opening of a gate in the replisome to allow the replisome to move past the ICL. Whether this is the gate between MCM2-MCM5, which would be unlocked by the loss of the GINS ^51^, or an alternative gate which can open independently of the GINS status ^52,53^ remains to be determined ^54^. Additionally, the translocase activity of FANCM could be required for moving the opened replisome past the ICL or modulating DNA structure once past the barrier. These are not exclusive possibilities.

In contrast to FANCM, DONSON has no enzymatic activity. Consequently, it may serve as a recruitment platform for factors that promote the stability of replication forks that encounter obstacles in early and mid S phase. For example, these might include enzymes such as SMARCAL1 and ZRANB3 that protect forks from collapse, and would provide a translocase activity, perhaps similar to FANCM ^55,56^. DONSON has also been demonstrated to be required for efficient activation of the ATR-dependent replication stress response ^30^.

Eu- and hetero-chromatin domains are not absolute, but subject to alteration during development, neoplasia, and aging ^35,57,58^. It will be of interest to determine the influence of these changes on the response to replication stress by DONSON and FANCM associated replisomes.

## Methods

### Data reporting

Statistical methods were not used for sample size determination. The experiments were not randomized, and the investigators were not blinded during experiments and data analysis.

### Materials

Dig-TMP was synthesized as described previously ^25^. The siRNA for DONSON and FANCM were purchased from Dharmacon. L-017453-02-0005, ON-TARGETplus Human DONSON siRNA SMART pool (GAAAUCAUCUUUACGGAAU, UGGACAAAGUACUUGA UAU, GAGAUGGGUGUGCAAGAUA, ACUUAGUCAAAUACCGUUA). L-021955-00-0005, ON-TARGETplus Human FANCM siRNA – SMART pool (GGGUA GAACUGGCCGUAAA, GAGAGGAACGUAUUUAUAA, AAACAGACAUCGCUGAAUU, GCAUGUAGCUAGGAAGUUU). Other reagents were Lipofectamine RNAiMAX (Invitrogen, 13778-150), Halt™ Protease and Phosphatase inhibitor cocktail (Thermo Scientific, 78446) and ATR inhibitor (VE821, Selleckchem, S8007).

### Cells, cell culture, transfection

Hela CCL-2 and RPE1 (ATCC) cells were maintained in DMEM (Gibco) supplemented with 10% fetal calf serum (Gibco), 100 U/mL penicillin, and 100 μg/mL streptomycin sulfate (Gibco). GFP-DONSON expressing HeLa cells ^30^ were cultured with L-Glutamine (Gibco), 200 ug/ml Hygromycin B (Invitrogen), and 5 ug/ml Blasticidin (Gibco). HeLa-Flp-In T-REx cells stably transfected with pcDNA5/FRT/TO-EGFP expressing EGFP or EGFP-DONSON were induced by incubation with 1 μg/ml Doxycycline for 48 hr. Cells derived from patient 9 with mutations in DONSON ^30^, stably transduced with pMSCV-vector only or pMSCV-DONSON, were grown in DMEM (Gibco) supplemented with 10% fetal calf serum (Gibco), L-Glutamine (Gibco), 100 U/mL penicillin, 100 μg/mL streptomycin sulfate (Gibco). All cells were routinely tested for mycoplasma (Lonza, LT07-701). To determine the effect of knock down of DONSON and FANCM in the DNA fiber assay, Hela cells were transfected with 10 nM siRNA (Dharmacon) using RNAiMAX (Invitrogen) on day 1 and day 2. Experiments were performed on day 4, ie.72 hr after siRNA transfection.

### Chromatin extraction and Immunoprecipitation

10^7^ cells were suspended in buffer A (10 mM HEPES at pH 7.9, 10 mM KCl, 1.5 mM MgCl2, 0.34 M sucrose, 10% glycerol, 1 mM DTT, 10 mM NaF, 1 mM sodium orthovanadate, 0.1% Triton X-100, with protease and phosphatase inhibitors) and incubated for 5 mins on ice. Nuclei were recovered by centrifugation at 1300 g for 4 min. The nuclear pellet was lysed in buffer B (3 mM EDTA, 0.2 mM EGTA, 1 mM DTT, protease and phosphatase inhibitors) for 10 min on ice, and then centrifuged at 1700 g for 4 min. Chromatin was resuspended in benzonase buffer (Sigma, E8263, 250 U/mL benzonase, 20 mM Tris-HCl at pH 8.0, 0.2 mM MgCl2, 2 mM NaCl, protease and phosphatase inhibitors and incubated at 4 °C overnight. Another 250 U/ml benzonase was added and the sample incubated for an additional 3 hrs. The sample was clarified by centrifugation and the supernatant adjusted to 200 mM NaCl, 50 mM Tris-HCl pH 7.4, 0.1% Tween 20.

For immunoprecipitation, soluble chromatin samples were precleaned with Dynabeads Protein G (Life Technologies) for 1h at room temperature. Then incubated with specific antibodies at 4 °C overnight. For sequential Co-IP, immunoprecipitations were performed using protein G magnetic beads (Pierce, 10% v/v), GFP Trap (Chromotek, gta-20). We performed each immunocapture twice, in order to clear the target complex. After capture with one antibody was completed the supernatant was incubated with the next antibody, and so on. All bead-antibody complexes were washed three times with PBS-T (phosphate buffered saline, .05% Tween-20. pH 7.5) and resuspended in SDS PAGE loading buffer. After heating for 10 min at 90 ℃, the proteins were analyzed by western blotting according to standard procedures.

### In situ Proximity Ligation Assay (PLA)

Cells were grown on Mattek glass bottomed plates followed by treatment with 5 μM Dig-TMP/UVA, 1.5 μM TMP/UVA, or UVA only. UVA exposure was in a Rayonet chamber at 3 J/cm^2^. After incubation with fresh medium for 60 min, cells were incubated with 0.1 % formaldehyde for 5 min and then treated twice with CSK-R buffer (10 mM PIPES, pH 7.0, 100 mM NaCl, 300 mM sucrose, 3 mM MgCl2, 0.5%Triton X-100, 300 μg/ml RNAse) and fixed in 4% formaldehyde in PBS (W/V) for 10 min at RT, followed by incubation in pre-cold methanol for 20 min at −20 °C. After washing with PBS cells were treated with 100 ug/ml RNAse for 30 min at 37 °C. *In situ* PLA was performed using the Duolink PLA kit (Sigma-Aldrich) according to the manufacturer’s instructions. Briefly, cells were blocked for 30 min at 37 °C and incubated with the respective primary antibodies (see reagent list) for 30 min at 37 °C. Following three times washing with PBST (phosphate buffered saline, 0.1% Tween), anti-Mouse PLUS and anti-Rabbit MINUS PLA probes were coupled to the primary antibodies for 1 h at 37 °C. After three times washing with Buffer A (0.01 M Tris, 0.15 M NaCl and 0.05% Tween 20) for 5 min, PLA probes were ligated for 30 min at 37 °C. After three times washing with Buffer A, amplification using Duolink In Situ Detection Reagents (Sigma) was performed at 37 °C for 100 min. After amplification, cells were washed for 5 min three times with Wash Buffer B (0.2 M Tris 0.1 M NaCl). Finally, they were coated with mounting medium containing DAPI (Prolong Gold, Invitrogen). Antibody specificity was confirmed by omitting one or another antibody. In some experiments, after completion of the PLA procedure, the stage of S phase was determined by immunostaining of cells with an antibody against PCNA conjugated with Alexa 647.

### PLA imaging and quantification

PLA plates were imaged on a Nikon TE2000 spinning disk confocal microscope, using a Plan Fluor ×60/1.25 numerical aperture oil objective. All images in an experiment were acquired with the same exposure parameters. Quantification was done on CellProfiler using the pipeline provided as Supplementary Information 2. Briefly, the pipeline performs the following steps: identify nuclei using the DAPI channel, filter to a maximum size the PLA foci, mask the foci image using the nuclei objects (PLA foci) to generate a visual representation of the foci counted for each cell, identify primary objects (PLA foci), establish a parent-child relationship between the foci (“children”) and nuclei (“parents”) in order to determine the number of foci per nucleus and export results as number of PLA foci per nucleus to a spreadsheet. The spreadsheets were compiled in Excel and exported to Graphpad Prism to generate the dot plots and determine if differences were statistically significant using the Mann-Whitney Rank sum test (NS: p>0.5, significant: p<0.001).

### Sequential PLA and 3D reconstruction

Mattek glass bottomed plates were marked on the growth surface with a diamond pen prior to plating cells in order to provide a reference for location of individual microscope fields ^59^. GFP-DONSON: pMCM2S108 PLA was performed as above, with Duolink Detection Reagent Green (Sigma, DUO92014) or Orange (DUO92007). Bright field images of the individual fields were obtained as well as the patterns of the PLA. The plates were then incubated with 6 M Guanidine: HCl in 5 % sucrose for 10 min at 40 ⁰C to strip the antibodies and reaction products, and then washed with PBST. The fields were inspected to ensure complete removal of signal after which the FANCM: pMCM2 S108 PLA was performed, with detection oligonucleotides linked to Duolink Detection Reagent Red (Sigma, DUO92013). The cells photographed after the first PLA were located and imaged again. 16 stacks covering 1.6 um were acquired of the first and second PLAs using Volocity software and exported as .OMETIFF. Both sets of images were converted into .ims to generate the 3D reconstructions on IMARIS (Bitplane) as follows. One of the sets (PLA2) was imported as a timepoint into the other set (PLA1). Next, a surface of each nuclei was created in the DAPI channel in order to track and correct for translational and rotational drift between the two PLA images. Each timepoint was then saved as the corresponding PLA, and the 4 channels were finally combined into one image to confirm correct alignment of nuclei (on the DAPI channel) and visualize the localization of both PLA signals on the same cell. A 3D reconstruction of one of such merged images is provided as Supplementary Movie 1.

### DNA Fiber Analysis

DNA fiber assays were performed as described previously ^25^. Briefly, cells were incubated with 6 μM Dig-TMP at 37 °C for 1 hr, followed by exposure to UVA light in a Rayonet chamber at 3 J/cm2 prior to incubation with 10 μM CldU for 20 min and then with 100 μM IdU for 20 min. Cells were trypsinized and suspended in PBS and approximately 200 cells placed on a glass microscope slide (Newcomer Glass) and 10 ul of lysis buffer (0.5% SDS in 200 mM Tris-HCl pH 7.5, 50 mM EDTA) added. DNA fibers were spread and fixed in 3:1 Methanol: Acetic acid, denatured with 2.5 M HCl for 1hr, neutralized in 0.4 M Tris-HCl pH 7.5 for 5 min, washed in PBS, and immunostained using anti-Dig, anti-BrdU primary and corresponding secondary antibodies. Antibodies and dilutions were rat anti-BrdU (CldU), 1:200; Dylight 647 goat anti-rat, 1:100; mouse anti-BrdU (IdU), 1:40; and Dylight 488 goat anti-mouse, 1:100 and Qdot 655 goat anti-mouse 1: 2,500. The slides were mounted in ProLong Gold Antifade Mounting medium. Images were acquired using a Zeiss Axiovert 200 M microscope at 63× magnification with the Axio Vision software packages (Zeiss). The quantum dot signal was imaged with a Qdot 655 filter.

### Analysis of early and late S phase cells

GFP-DONSON expressing cells were treated with 1.5 μM TMP/UVA and after 1 hr were trypsinized and suspended in DMEM with 10 % fetal calf serum and incubated with 16 μM Hoechst for 30 min at room temperature. The cells were centrifuged, washed with sorting buffer (HEPES pH 7.0, 1 mM EDTA, and 5 % fetal calf serum), and then suspended in 1 ml of sorting buffer supplemented with 1 mM N-acetyl cysteine. The cells were then resolved by flow cytometry and early and late S phase fractions harvested. Cells from each fraction were attached to slides by centrifugation (Cytospin), fixed with 0.1 % formaldehyde and PLA between GFP-DONSON: pMCM2S108 or FANCM: pMCM2S108 performed.

### Chromatin Immunoprecipitation (CHIP) for DNA analysis

Cells were crosslinked with 1% formaldehyde in culture media for 8 min, followed by quenching the formaldehyde with 0.1 M glycine. Cells were washed with PBS, harvested by scraping, then suspended in lysis buffer (0.5% SDS, 10 mM EDTA, 50 mM Tris-HCL pH 8.0) supplemented with protease and phosphatase inhibitors. Lysates were sonicated in a 4 ⁰C water bath ultrasonicator (Bioruptor, Diagenode). The time of sonication was adjusted to yield short DNA fragments <500 bp (total 8 minutes, 30 seconds sonication, then cool 30 seconds). In some experiments the time was adjusted to yield longer DNA fragments of 500-5000 bp (2 x 30 seconds with a 30 second cooling period). Diluted lysates were incubated overnight at 4 °C with antibodies as indicated. Immunoprecipitations were performed using Protein G magnetic beads (Pierce, 10% v/v), or GFP Trap (Chromotek, gta-20). Bead bound complexes were washed with low salt immune complex buffer (0.1% SDS, 1% Triton x-100, 2 mM EDTA, 20 mM Tris-HCl pH 8.0, 150 mM NaCl), high salt immune complex buffer (0.1% SDS, 1% Triton x-100, 2 mM EDTA, 20 mM Tris-HCl pH 8.0, 500 mM NaCl), LiCl immune complex buffer (0.25 M LiCl, 1% NP-40, 1% mM EDTA, 10 mM Tris-HCl pH 8.0) and TE buffer (10 mM Tris-HCl, 1 mM EDTA pH 8.0). DNA was eluted in elution buffer (1% SDS, 0.2 M NaCl) with Proteinase K (100 μg/ml) overnight at 65 °C. Eluted DNA was purified with DNA Clean & Concentrator PCR purification Kit (ZYMO Research, D4033) according to the manufacturer instructions.

### Dot blot analysis

The DNA was denatured using 0.5 M NaOH and 1.5 M NaCl and equal amounts were loaded onto a Hybond N + nitrocellulose membrane (GE Biosciences) using the Bio-Dot apparatus (Bio-Rad). Membranes were washed once with denaturing buffer and wash buffer (3× SSC), followed by UV-crosslinking (UV Stratalinker 1800, Stratagene) and blocking with 5× Denhardt’s solution (Thermo Scientific) for 1 h at 37 °C. Hybridization with Alu-Biotin (5′ Biotin- GGCCGGGCGCGGTGGCTCACGCCTGTAATCCCAGCA), Satellite III (5′ Biotin- TCCACTCGGGTTGATT) or LINE-1 (5′ Biotin- GACTTCAAACTATACTACAAGGCTACAGTAACC) probes was performed at 37 °C overnight. Chemiluminescent Nucleic Acid Detection Module Kit (Thermo Scientific, 89880) was used for signal detection and images were acquired using ChemiDox XRS with Image Lab software (Bio-Rad).

### Western blotting

For a full list of antibodies, see reporting summary. The samples were prepared in NuPAGE Sample Buffer (Invitrogen). Then proteins were separated by electrophoresis in 4%–12% Bis-Tris Protein Gels and transferred to polyvinylidene difluoride membrane (Thermo Scientific). The membranes were blocked in 5% dry milk in 0.1% Tween-20 in PBS and detected with the indicated antibodies. After incubation with horseradish peroxidase (HRP)-conjugated secondary antibodies (BIO-RAD), proteins were visualized using ECL detection reagents (GE Healthcare). Uncropped gel images for western blots are available in Supplementary information 3.

### Statistics and reproducibility

Statistical significance of PLA experiments was analyzed using the Mann-Whitney Rank sum test. Fiber patterns and immunoblotting were analyzed using a two-sided unpaired t-test and the exact p-values are given in each case. For both tests: Significant: p<0.001, NS (not significant): p>0.05. All experiments were performed at least twice and the number of biological replicates (n) is reported in each figure legend.

### CHIP-SEQ

Immunoprecipitation of sonicated chromatin was performed as described above.

### DNA sequencing

For DNA sequencing, Illumina sequencing adapters with a T-overhang were ligated to the precipitated ChIP DNA fragments or the input DNA, with a corresponding A-overhang, to construct a sequencing library according to the manufacturer’s protocol (Illumina, San Diego, CA). The fragments were purified using a magnetic bead protocol and eighteen cycles of PCR amplification were performed to enrich for fragments with an adapter on both ends. The products were purified again with size selection (approximately 200–600 bases) using a dual bead selection protocol with SPRIselect Beads (Beckman Coulter, Brea, CA). These libraries were sequenced on an Illumina Hi-Seq 2500 sequencer using on-board cluster generation on a rapid run paired end flow cell for 75 × 75 cycles (DONSON) and single end of 75 bp for 75 cycles (FANCM). Real-time analysis was performed using RTA v1.18.66.3 and base-calling was performed using bcl2fastq v2.18.0.12.

### Chip-Seq, RT and Hi-C data

The log2 ratio between FANCM or GFP_DONSON ChIP-seq and the Input was computed using Deeptools BigWigCompare of the corresponding RPKM normalized BigWig files. All datasets generated in this study are deposited in the National Center for Biotechnology Information Gene Expression Omnibus (GEO) database (https://www.ncbi.nlm.nih.gov/geo/; GEO series XXXXXXX). H3K9me3 and H3K4me3 ChIP-seq data from Hela cells was downloaded from *GEO:* GSM2514495 and GSM3398459, respectively. Replication Timing data for Hela S3 cells was downloaded from the Replication Domain database, curated by the Gilbert laboratory(https://www2.replicationdomain.com/#): RT_HeLaS3_CervicalCarcinoma_Int 2355 8071_hg38. Hi-C data for HeLa cells was downloaded from NCBI, dbGaP phs000640.v8.p1. [https://doi.org/10.1016/j.cell.2014.11.021] The eigenvector, used to delineate compartments in Hi-C data at coarse resolution, was calculated as the first principal component of the Pearson’s matrix using Juicer (-p KR, BP 50,000). UCSC liftover was used to convert hg19 to hg38 genome coordinates. Fragile sites mapping coordinates were downloaded from HumCFS: a database of Human chromosomal fragile sites. https://webs.iiitd.edu.in/raghava/humcfs/download.html.

### RT scores for FANCM and GFP-DONSON enriched genomic windows

The genome was divided into 50 Kb windows and the mean RT score for each window was calculated. The genomic regions enriched in FANCM and GFP-DONSON were selected as those 50 Kb windows in each ChIP with a [log2(ChIP/Input) > 0]. Their corresponding RT scores were determined, and their distribution of RT scores mapped as violin plots. The box plot inside the violins represent the median and the interquartile range. Randomly selected genomic regions, with number and size of the genomic windows matching each sample, were used as controls. All of the randomized samples have equivalent distributions. In the case of the eigenvector, we used 50 Kb genomic regions with a value > 0 for the A compartment, and < 0 for the B compartment. As expected, the distribution of RT scores is heavily biased towards early replication for the A compartment, and late replication in the B compartment.

### Chip-seq coverage of RT quantiles

The genome was divided into 50 Kb windows and the mean RT score for each window was calculated. The coverage of each ChIP BAM file per genomic window was computed and the counts converted to TPM (tags per million). The RT quantiles were calculated (n=25). The chip-seq coverage was displayed in TPM for each of the 25 RT quantiles ordered from Late to Early.

### Chip-seq coverage of Hi-C eigen vector quantiles

The eigenvector, used to delineate compartments in Hi-C data at coarse resolution, was calculated as the first principal component of the Pearson’s matrix using Juicer (-p KR, BP 50,000). Chip-seq coverage of eigenvector quantiles was calculated as follows: the genome was divided into 50 Kb windows and the mean eigen vector score for each window calculated. We then computed the coverage of each Chip-seq BAM file for each genomic window, converted counts to TPM (tags per million) and calculated Hi-C compartment eigenvector quantiles (n=25). ChIP-seq coverage was displayed in TPM for each of the 25 eigen vector quantiles ordered from B to A.

### Reporting summary

Further information on research design is available in the Nature Research Reporting Summary linked to this paper.

### Data availability

All datasets for this study are available from the corresponding author on request. Code will be made available upon request.

## Acknowledgements

This research was supported, in part, by the Intramural Research Program of the NIH, National Institute on Aging, United States (<u>Z01-AG000746-08</u>). GSM and GSS received funding from the CR-UK Program grant (C17183/A23303) and JJR was supported by the University of Birmingham. We thank Cuong Nguyen and Tonya Wallace of the NIA Flow Cytometry Unit for expert support and guidance, Supriyo De and William Wood for DNA sequencing and data extraction, Dr. Florencia Pratto for data analysis, and Dr. Yie Liu, Dr. Rafael D. Camerini-Otero, and Dr. Weidong Wang for helpful suggestions and discussions.

## Author Contributions

J.Z., R.J., M.A.B., D.P., M.M.S designed and performed experiments, analyzed data, and prepared figures. G.S.M., J.J. R. constructed cell lines, A.P.J. generated antibody against DONSON, G.S.S., supervised and contributed to the writing of the manuscript, J.Z and M.M.S. conceived the study, designed experiments, analyzed data, J.Z., M.A.B., and M.M.S. wrote the manuscript.

## Corresponding author

Michael M. Seidman

## Figure Legends

**Supplementary Figure 1:**
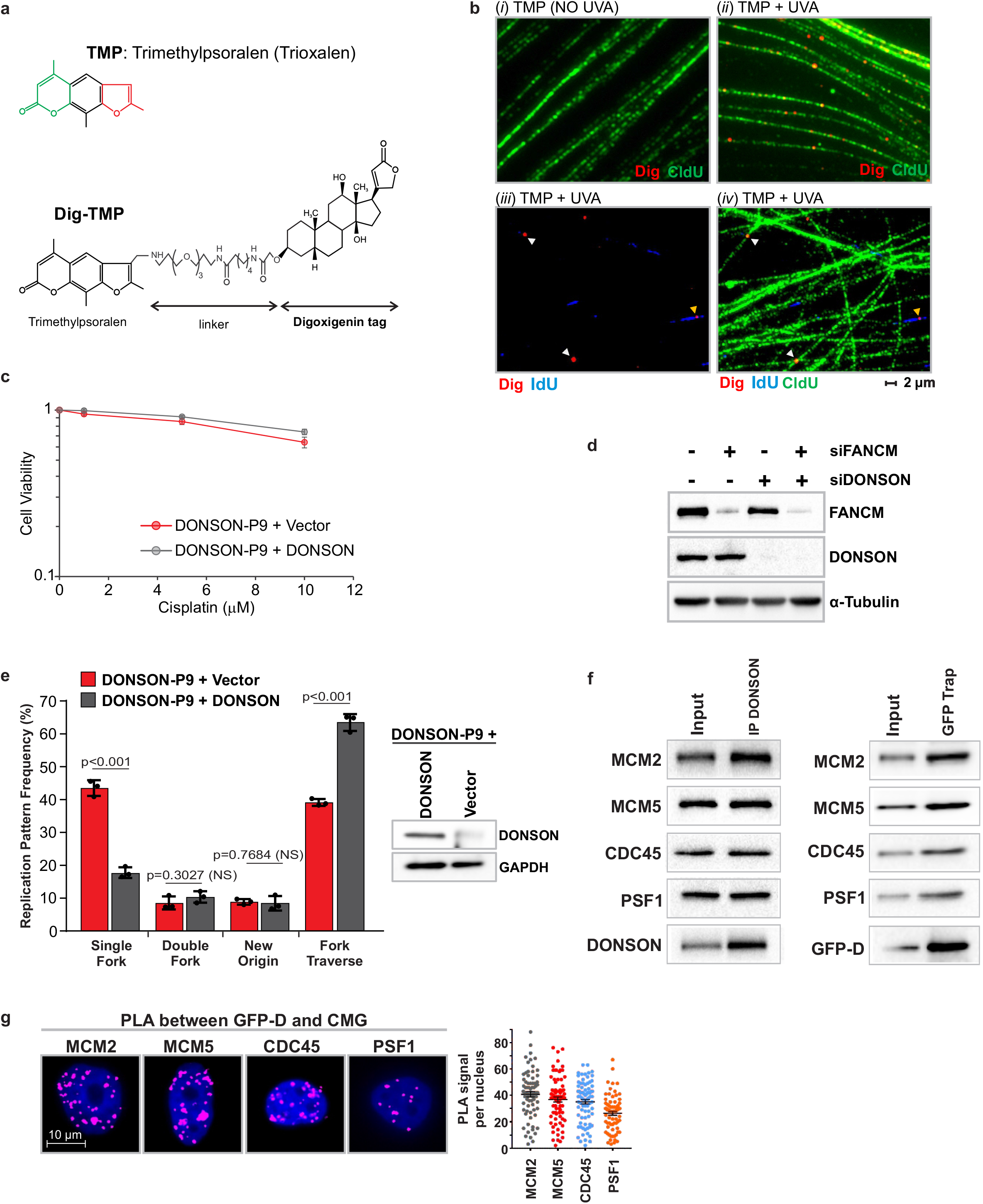
DONSON contributes to replication traverse of ICLs. **a**. Structure of Digoxigenin-tagged trimethyl psoralen. **b**. Fibers from cells exposed to Dig-TMP/UVA. i. Cells were incubated with CldU 24 hrs, then with Dig-TMP. The UVA exposure, required for crosslinking, was omitted. Fibers were displayed by immunofluorescence (green), and a primary antibody against Dig and a secondary tagged with Q-dot 655 (red). The absence of Q-dot signals reflects the absence of covalently bound Dig tagged ICLs. **ii**. Cells were incubated with 20 μM Dig/TMP (a higher concentration than in replication experiments) and exposed to UVA. Note the presence of numerous ICLs on the fibers. **iii**. Cells were incubated with CldU for 24 hrs, treated with 6 μM Dig-TMP/UVA, then incubated with IdU for 30 minutes. The IdU and Dig signals are shown, and an encounter with an ICL denoted (yellow arrow). Dig signals (white arrows) not associated with an IdU tract. **iv.** Display of CldU and IdU and Dig in the field shown in iii. Dig-TMP signals are on fibers labeled by CldU, although they may not be associated with an IdU tract (yellow arrow). **c**. DONSON does not contribute to survival of cells exposed to Cisplatin. Patient derived cells DONSON-P9 were complemented with either wild type DONSON or vector only. **d**. Knockdown efficiency of siRNAagainst FANCM, DONSON, or FANCM/DONSON in HeLa cells. **e**. Single fork stalling at ICLs is increased in patient derived cells DONSON-P9. The replication traverse assay was performed in patient derived cells complemented with the vector alone or wild type DONSON. **f**. Immunoprecipitation of either endogenous DONSON or GFP-DONSON demonstrates association with replisome proteins. **g**. The proximity of GFP-DONSON and replisome proteins demonstrated by PLA. Scored nuclei: GFP-DONSON and MCM2= 76, GFP-DONSON and MCM5= 79, GFP-DONSON and CDC45= 80, GFP-DONSON and PSF1= 76, from 3 biological replicates. A two-sided unpaired t-test was used to calculate p-values for replication pattern frequency experiments. Data are mean ± s.d. Mann-Whitney Rank sum test was used to calculate p-values for PLAexperiments. Data are mean ± s.e.m. NS: not significant (p>0.05).

**Supplementary Figure 2:**
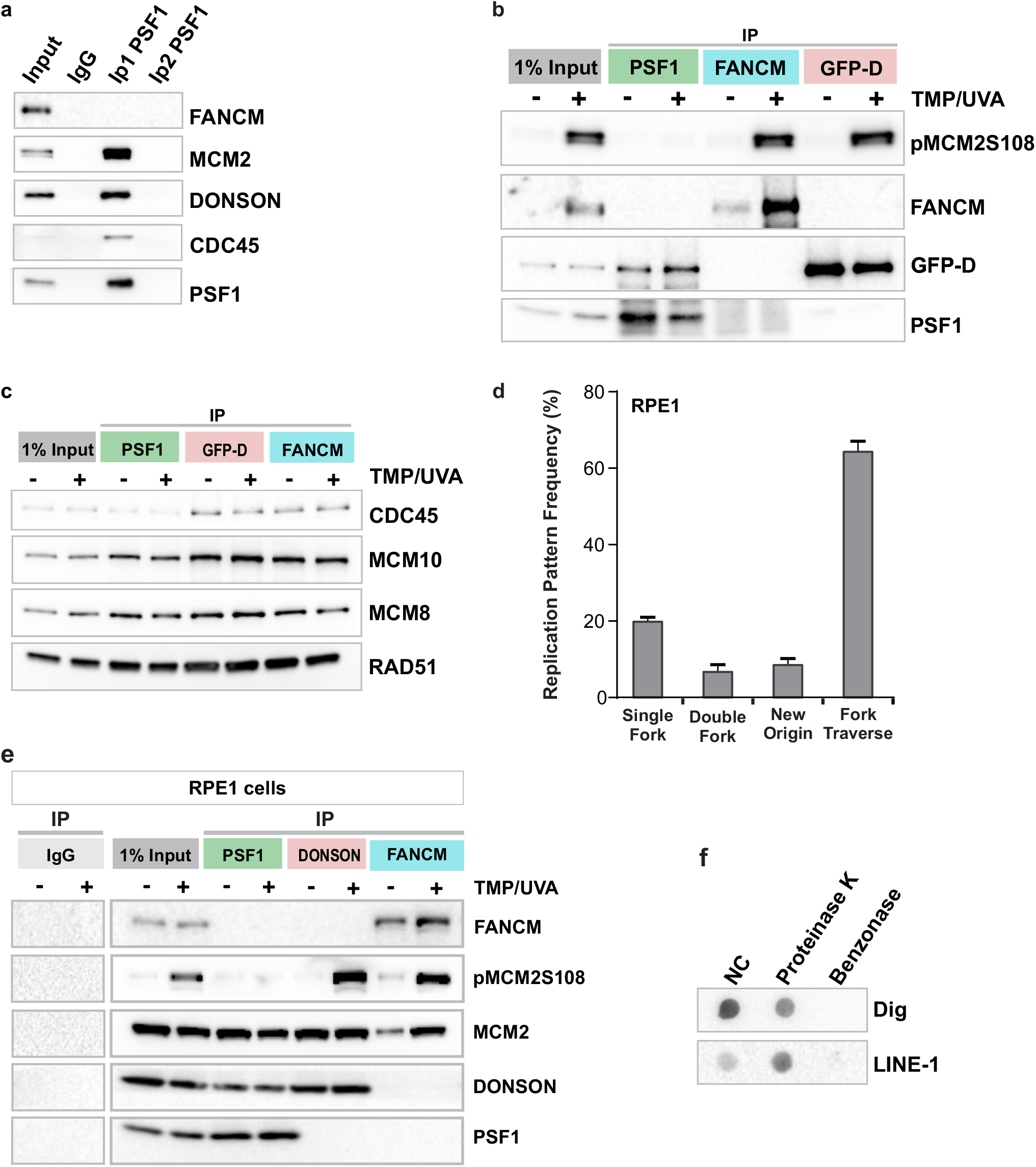
DONSON and FANCM are on different replisomes. **a.** Efficacy of IP against the GINS protein PSF1. Chromatin proteins were incubated with antibody against PSF1 and the precipitate removed. No PSF1 was recovered when the supernatant was challenged again with the same antibody. Representative blot (n = 2). **b**. Reversal of the order of the sequential IP of replisome components does not change the results. IP against FANCM preceded IP against GFP-DONSON. Representative blot (n = 2). **c**. Proteins common to each replisome complex. Representative blot (n = 3). **d.** Replication patterns in RPE1 cells containing ICLs are the same as in other cells. **e**. DONSON and FANCM are on different replisomes in RPE1 cells. **f**. Dig-TMP in sonicated chromatin is associated with DNA. To verify the covalent linkage of Dig-ICL with DNA the sonicated chromatin used for sequential IP was digested with either benzonase or proteinase K. The samples were then examined by dot blot for the Dig tag on the ICLs. Representative blot (n = 3).

**Supplementary Figure 3:**
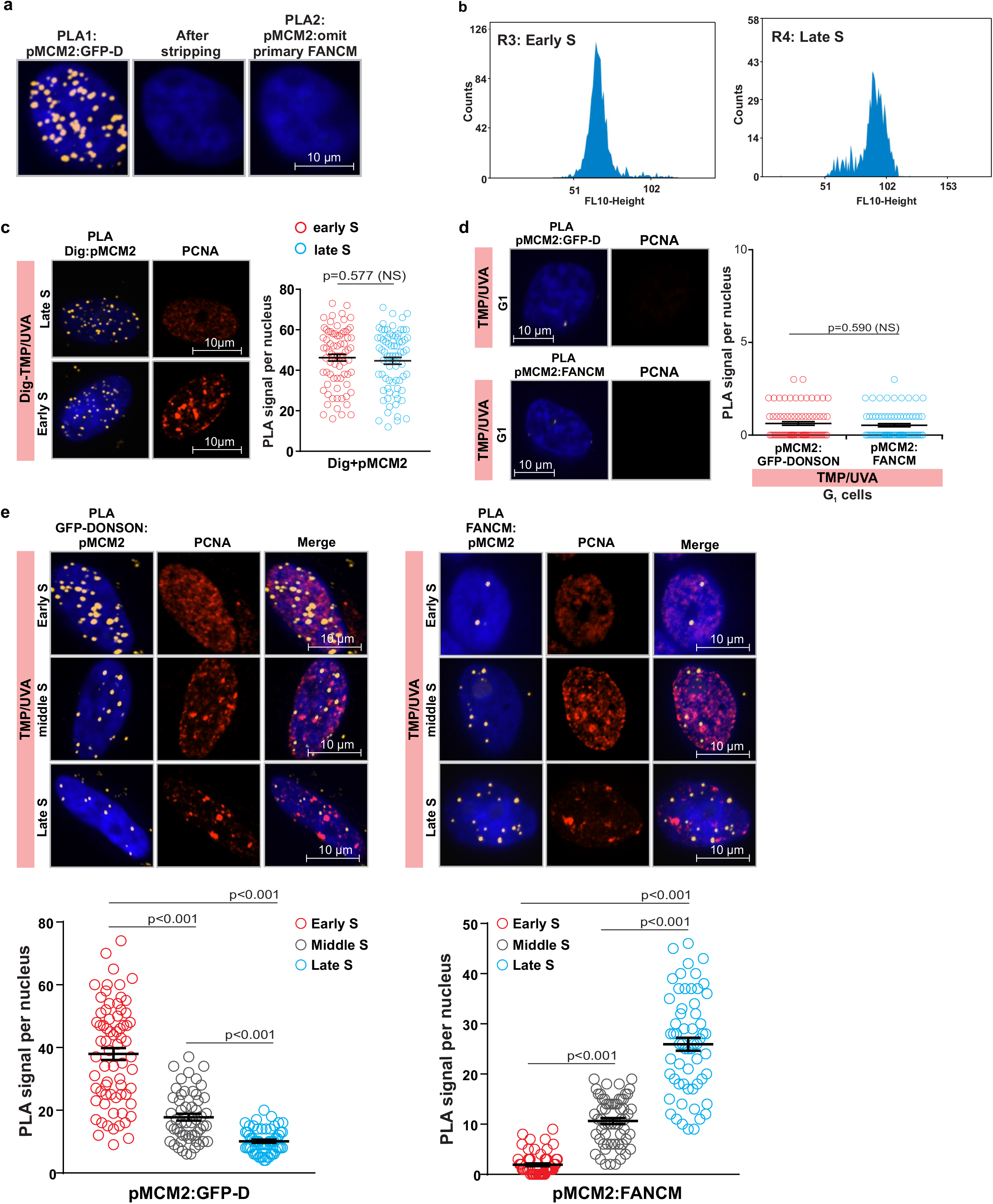
DONSON and FANCM replisomes are in different locations and active at different times of S phase. **a**. Antibody omission control for sequential PLA. Accurate interpretation of sequential PLA (Methods) requires complete removal of the antibodies and reaction products of the first reaction, in order to avoid compromising the second reaction. The GFP-DONSON: pMCM2S108 PLA was performed. After image acquisition the cells were stripped of the components and products of the PLA and a second PLA was performed without the addition of the antibody against FANCM. The absence of signal demonstrates the efficacy of the stripping procedure. **b**. Sorted early and late S phase cells are not cross contaminated. Early and late S phase cells were recovered and re-analyzed. **c**. The frequency of replisome encounters with ICLs is similar in early and late S phase. The PLA between pMCM2 and the Dig tag on the ICLs shows equivalent frequencies in early and late S phase. **d**. Specificity test of DONSON and FANCM PLA with pMCM2S108. There can be no replisome encounters with ICLs in G1 phase cells. The PLA between GFP-DONSON or FANCM and pMCM2S108 was performed as a test of antibody and assay specificity. Scored nuclei of PLA between GFP-D and pMCM2S108= 75, FANCM and pMCM2S108= 75 from 3 biological replicates. **e**. The distinction between DONSON: pMCM2 replisomes and FANCM: pMCM2 replisomes in early and late S phase is lost in mid S phase cells. Scored nuclei: GFP-D and pMCM2S108, early S phase= 70, middle S phase= 53, late S phase= 59; FANCM and pMCM2S108, early S phase= 64, middle S phase= 61, late S phase= 60, from 3 biological replicates. Mann-Whitney Rank sum test was used to calculate the p-value for PLA experiments. Data are mean ± s.e.m. NS, not significant: p>0.05.

**Supplementary Figure 4:**
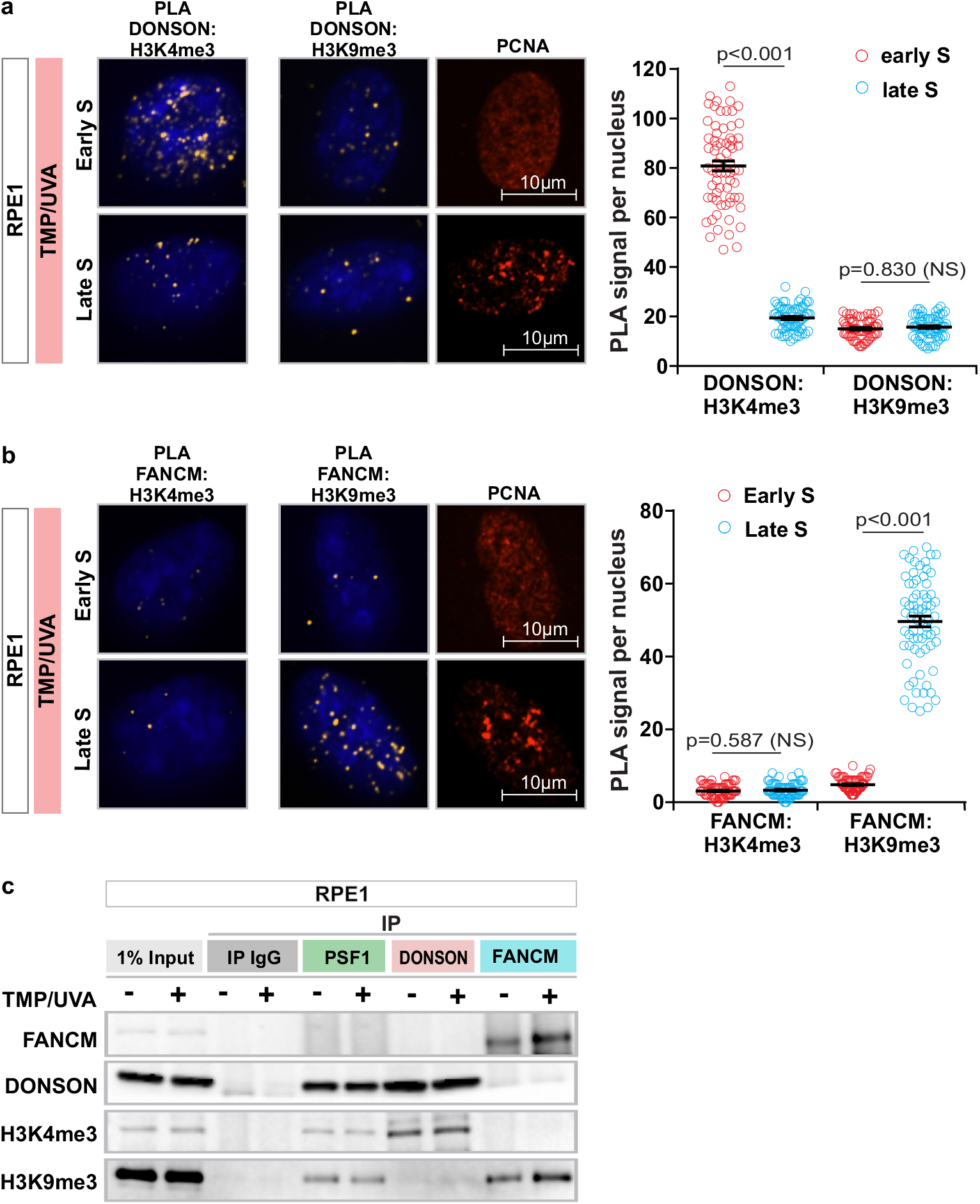
Association of DONSON and FANCM with H3K4m3 and H3K9me3 in RPE1 cells. RPE1 cells were treated with TMP/UVA and PLA between endogenous DONSON or FANCM and H3K4me3 or H3K9me3 performed. Signals were quantitated in early or late S phase cells. **a.** The association of DONSON with H3K4me3 is greater in early S phase cells than in late S phase. The interaction of DONSON with H3K9me3 is low in both early and late S phase cells. Scored nuclei: DONSON and H3K4me3, early S phase= 62, late S phase= 64; DONSON and H3K9me3, early S phase= 63, late S phase= 60, from 3 biological replicates. **b**. The association of FANCM with H3K4me3 is low in both early and late S phase, while that with H3K9me3 is much stronger in late than in early S phase. Scored nuclei: FANCM and H3K4me3, early S phase= 69, late S phase= 70; DONSON and H3K9me3, early S phase= 73, late S phase= 70, from 3 biological replicates. **c**. Cells were exposed to UVA or TMP/UVA. Sequential IP reveals greater association of DONSON with H3K4me3 than H3K9me3, and greater association of FANCM with H3K9me3 than H3K4me3. Mann-Whitney Rank sum test was used to calculate the p-value for PLA experiments. Data are mean ± s.e.m. NS, not significant: p>0.05.

**Supplementary Figure 5:**
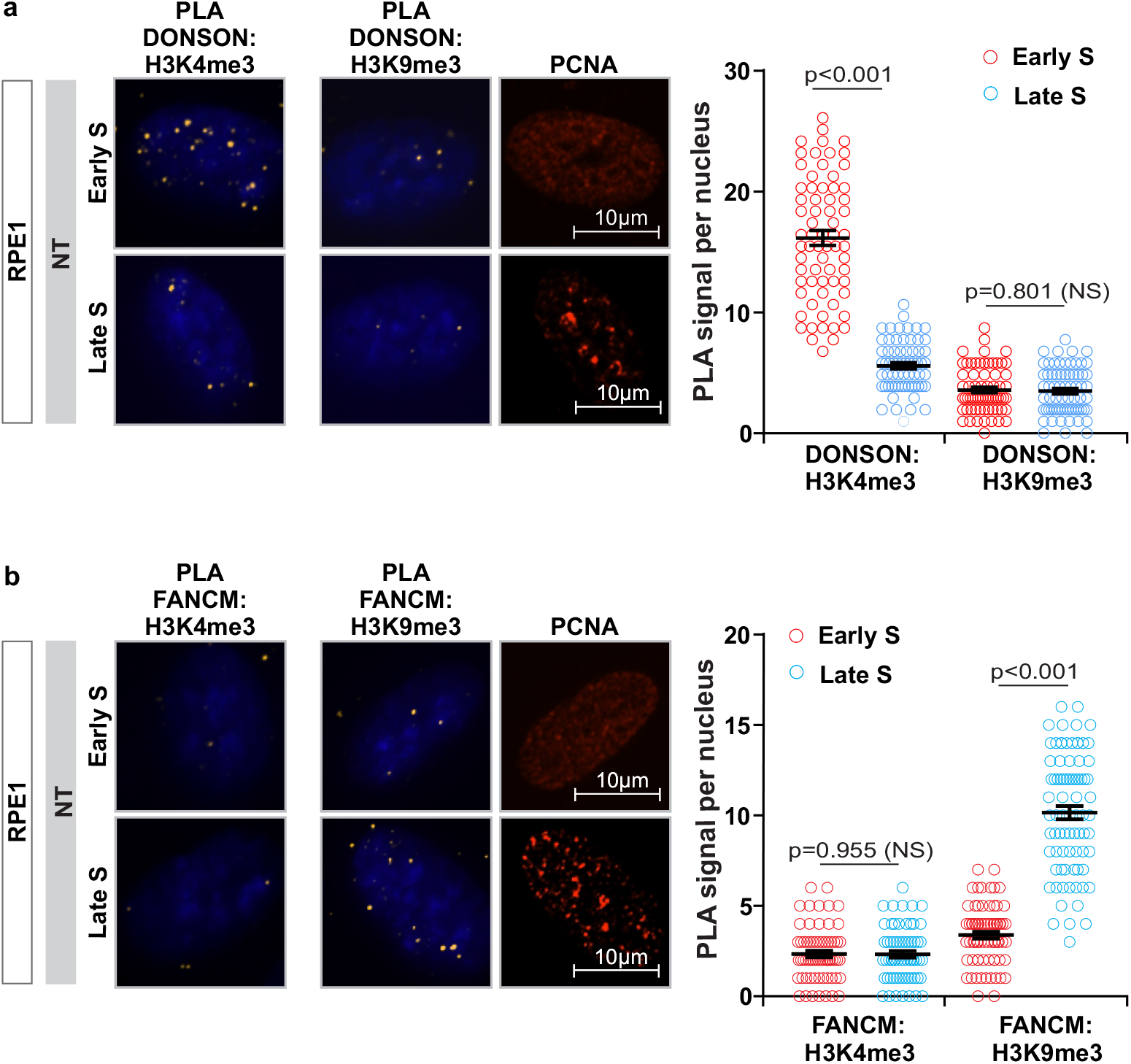
Interactions between DONSON or FANCM and H3K4me3 or H3K9me3 in untreated early and late S phase RPE1 cells. **a.** PLA between endogenous DONSON and H3K4me3 or H3K9me3. The association of DONSON with H3K4me3 is greater in early S phase cells than in late S phase. The interaction of DONSON with H3K9me3 is low in both early and late S phase cells. Scored nuclei: DONSON and H3K4me3, early S phase= 68, late S phase= 65; DONSON and H3K9me3, early S phase= 64, late S phase= 68, from 3 biological replicates. **b**. PLA between FANCM and H3K4me3 or H3K9me3. The association of FANCM with H3K4me3 is low in both early and late S phase, while that with H3K9me3 is much stronger in late than in early S phase. Scored nuclei: FANCM and H3K4me3, early S phase= 67, late S phase= 65; DONSON and H3K9me3, early S phase= 73, late S phase= 80, from 3 biological replicates. Mann-Whitney Rank sum test was used to calculate the p-value for PLA experiments. Data are mean ± s.e.m. NS, not significant: p>0.05.

**Supplementary Figure 6:**
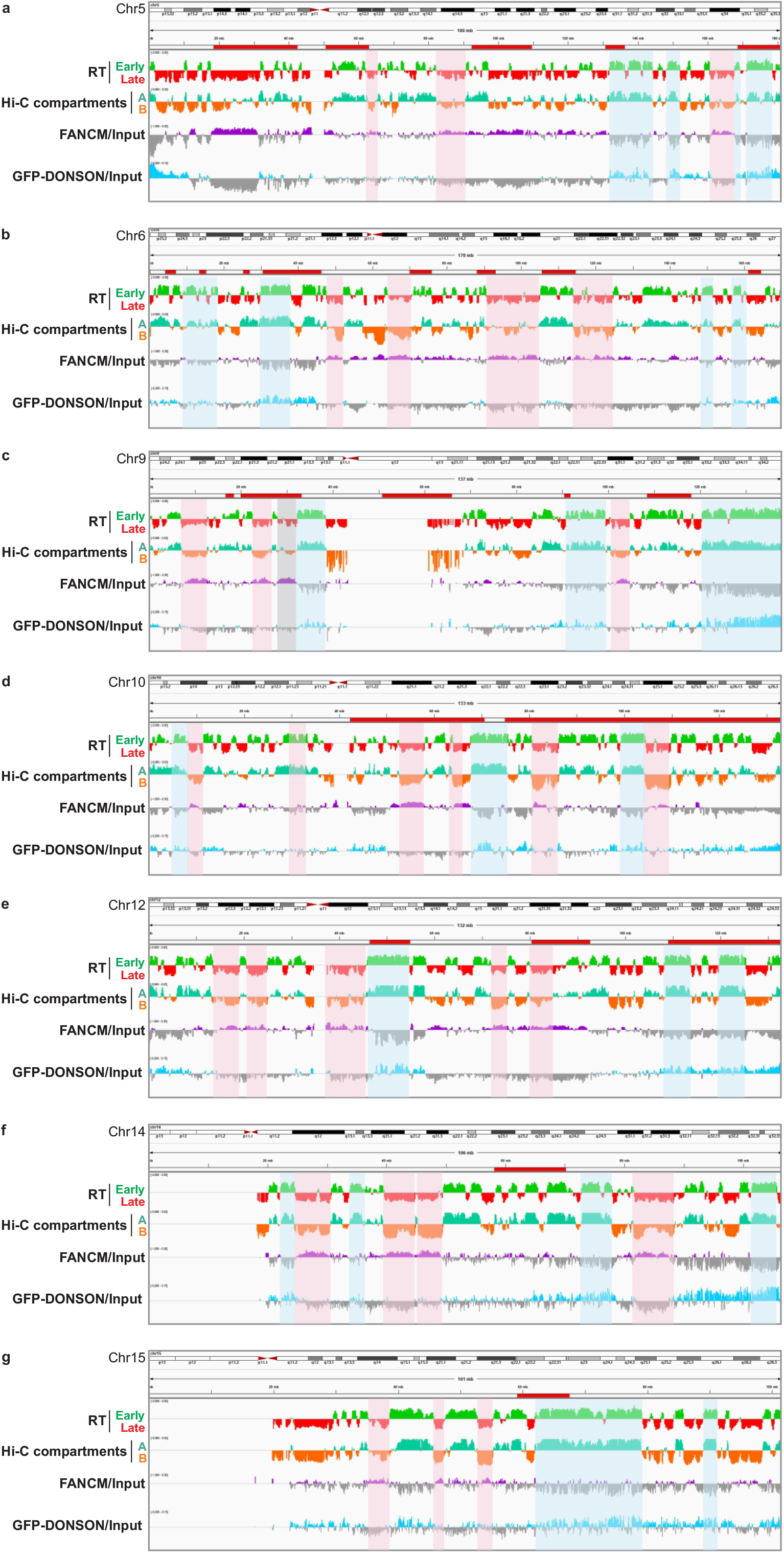
CHIP-seq distribution profiles for selected chromosomes. Correlations between FANCM and late replicating regions and chromatin compartment B are indicated in pink. Correlations of DONSON with early replicating regions and chromatin compartment A are indicated in blue. **a.** Chr 5. **b.** Chr 6. **c**. Chr 9. **d.** Chr 10. **e.** Chr 12. **f**. Chr 14. **g**. Chr 15.

